# End-to-end protein-ligand complex structure generation with diffusion-based generative models

**DOI:** 10.1101/2022.12.20.521309

**Authors:** Shuya Nakata, Yoshiharu Mori, Shigenori Tanaka

## Abstract

Three-dimensional structures of protein-ligand complexes provide valuable insights into their interactions and are crucial for molecular biological studies and drug design. However, their high-dimensional and multimodal nature hinders end-to-end modeling, and earlier approaches depend inherently on existing protein structures. To overcome these limitations and expand the range of complexes that can be accurately modeled, it is necessary to develop efficient end-to-end methods. We introduce an equivariant diffusion-based generative model that learns the joint distribution of ligand and protein conformations conditioned on the molecular graph of a ligand and the sequence representation of a protein extracted from a pre-trained protein language model. Benchmark results show that this protein structure-free model is capable of generating diverse structures of protein-ligand complexes, including those with correct binding poses. Further analyses indicate that the proposed end-to-end approach is particularly effective when the ligand-bound protein structure is not available. The present results demonstrate the effectiveness and generative capability of our end-to-end complex structure modeling framework with diffusion-based generative models. We suppose that this framework will lead to better modeling of protein-ligand complexes, and we expect further improvements and wide applications.

## Background

Molecular interactions between proteins and small molecule ligands are fundamental to biological processes, and three-dimensional structures of molecular complexes provide direct insights into their interactions associated with functions [1]. Since experimental structure determination is costly and often challenging, many computational methods have been developed for cheaper and faster modeling. Previous approaches predominantly employ the molecular docking methodology [2–8] that predicts preferred conformations of a ligand in a protein binding site. Despite its success in drug discovery and other applications, correct sampling of binding poses can be limited by poor modeling of protein flexibility [9–11]. Although numerous techniques [12–14] have been proposed to address this problem, they require manual setting with special attention or computationally expensive simulations for conformational sampling. Therefore, modeling the structure of protein-ligand complexes remains a major challenge, especially when the protein structure is flexible or unknown.

Advances in bioinformatics and deep learning have enabled accurate protein structure prediction using multiple sequence alignments (MSAs) and structural templates [15–17]. Subsequent studies have proposed single-sequence structure prediction methods utilizing protein language model (PLM) representations [18–20]. These methods have provided highly accurate structure predictions for proteins that have not been structurally characterized before, opening up new research possibilities. Nevertheless, an end-to-end structure prediction method for protein-ligand complexes that explicitly accounts for ligand features has not yet been established. While using protein structure predictions with docking is a promising approach, current prediction methods lack a principled way to provide diverse models, and obtaining a relevant model suitable for ligand docking is not always possible.

Deep generative modeling [21, 22] is a powerful approach to model high-dimensional and multi-modal data distributions and generate samples efficiently. In particular, diffusion-based generative models [23–26] have demonstrated a capacity for high-quality synthesis in many domains, including conformation generation of small molecules [27–30] and proteins [31–33]. A generative model that learns the joint distribution of protein and ligand conformations would enable principled sampling of diverse conformations and provide insights into their ensemble properties.

Recently, several methods have been proposed to apply diffusion-based generative models to protein-ligand complexes [34–36]. For instance, DiffDock [35] modeled the conformation of a ligand relative to a given protein with a diffusion-based generative model. The authors reported significant performance gains over existing methods on the PDBbind [37] benchmark dataset and highlighted the critical issues with regression-based frameworks [38, 39]. In particular, NeuralPLexer [34], similar work to ours, proposed to model the structure of protein-ligand complexes with a hierarchical diffusion model. However, this method still depended on protein backbone templates, and it is unclear whether diffusion models are applicable without structural inputs.

In this work, we propose an end-to-end framework to generate ensembles of protein-ligand complex structures by modeling their probability distributions with equivariant diffusion-based generative models (Figure 1). By incorporating the essence of the state-of-the-art protein structure prediction methods, the proposed framework can generate diverse structures of protein-ligand complexes without depending on existing protein structures, thus providing a novel and efficient method.

**Figure 1:**
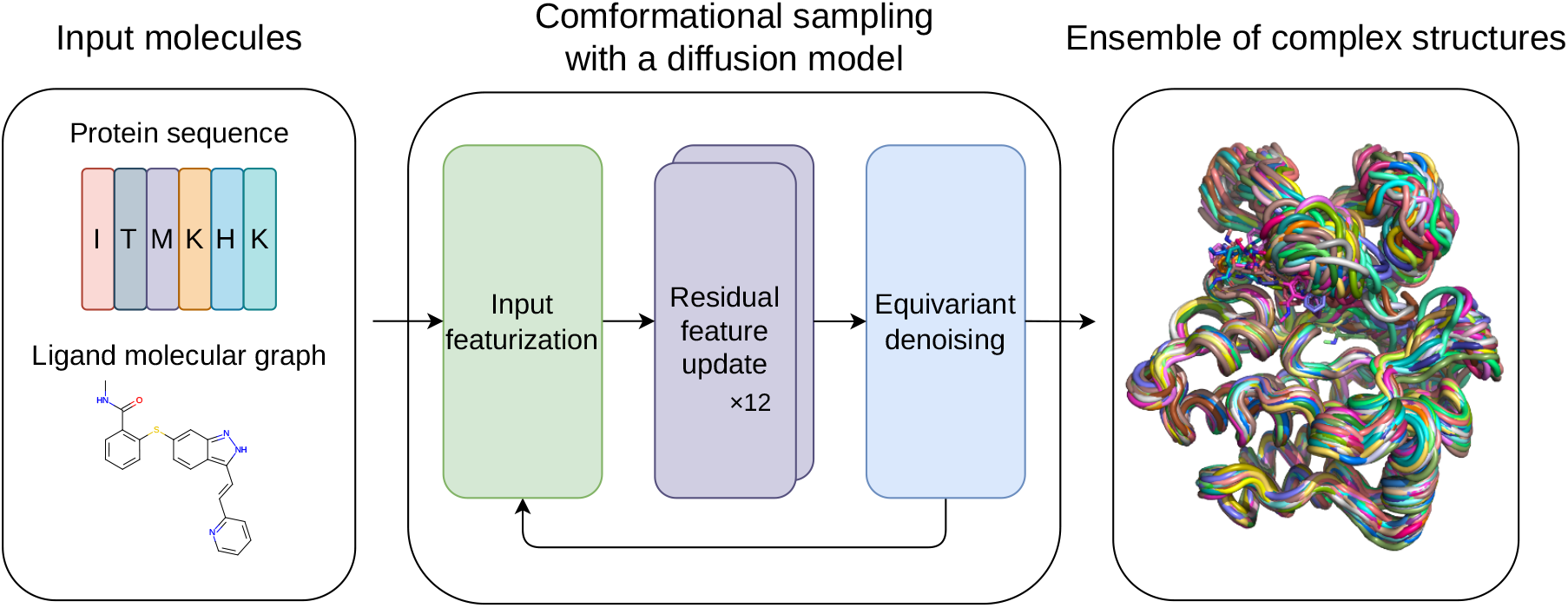
Overview of the proposed framework. For inputs, protein amino acid sequence and ligand molecular graph are employed. The conformational sampling process involves the iterative application of input featurization, residual feature update, and equivariant denoising to generate an ensemble of complex structures.

## Results

### Generative modeling of protein-ligand complex structures

We model the joint distribution of 3D coordinates of protein C_*α*_ and ligand non-hydrogen atoms with a variant of equivariant diffusion-based generative model [29]. The model uses a protein amino acid sequence and a ligand molecular graph as input and iteratively refines the 3D coordinates of the molecules starting from random noises to generate statistically independent structures.

We used protein-ligand complex structures from the PDBbind [37] database, a collection of biomolecular complexes deposited in the Protein Data Bank (PDB) [40], for training. We employed the time-based split proposed in EquiBind [38], where structures released in 2019 or later were used for evaluation, and any ligand overlap with the test set was removed for training and validation. For computational cost reasons, we only used structures with the number of modeled atoms (the sum of the numbers of protein residues and ligand non-hydrogen atoms) less than or equal to 384. This resulted in 9430 structures for training, 552 for validation, and 207 for evaluation. We used the Adam [41] optimizer with the base learning rate 4 × 10^−4^, *β*_1_ = 0.9, *β*_2_ = 0.999, *ϵ* = 10^−8^ and linearly increased the learning rate over the first 1000 optimization steps. We trained our model for around 150 epochs using a mini-batch size of 24. For evaluation, we used an exponential moving average of our parameters with the best validation loss, calculated with a decay rate of 0.999.

Though we designed our model to be protein structure-free, we also trained another version of the model that accepts protein backbone templates as input for reference. We denote the original protein structure-free model as DPL (Diffusion model for Protein-Ligand complexes) and this protein structure-dependent version as DPL+S.

### Benchmark on the PDBbind test set

To assess the generative capability of our models in terms of the reproducibility of the experimentally observed protein conformations and ligand binding poses, we sampled 64 structures for each complex in the PDBbind test set and compared them with the PDB-registered structures. We used TM-align [42] for structural alignment with experimental structures and used the resulting TM-score to evaluate protein conformations. For the evaluation of ligand binding poses, we calculated the heavy-atom root mean square deviation between generated and experimental ligands (L-rms) after aligning the proteins. As a baseline, molecular docking methods GNINA [7] and AutoDock Vina 1.2.0 [8] were used in blind self-docking settings with default parameters, except we increased exhaustiveness from 8 to 64 and num modes to 64.

Figure 2A shows the median modeling accuracies for each complex in the PDBbind test set. From this data, we can see that our models successfully reproduced the experimentally observed protein conformations and ligand binding poses for a significant number of complexes. Even though DPL did not use protein structures as inputs, it was able to sample a complex structure in which the protein conformation was close to the experimental one (TM-score *>* 0.9) for more than 70 % of the complexes in the test set (Figure 2B). Concerning the binding pose accuracies, our models achieved accuracy comparable to or better than baseline methods, especially in the range of practical importance, where the threshold is less than 2 Å (Figure 2C). Interestingly, the difference in binding pose accuracy between the two models DPL and DPL+S was insignificant. This is likely attributed to the fact that, as inferred from Figure 2A, generating the protein structure is almost always feasible if the model understands the complex well enough to reproduce the correct binding pose. Figure 2D shows how the performances change with the number of generative samples. Because our models generated diverse structures, a reasonable number of samples was needed to reproduce the experimental binding poses, while the performance approximately saturated after a few dozen samples. The relationships between the binding pose accuracy and the number of ligand rotatable bonds and that of related structures in the training set are shown in Figures 2E and F, respectively. Although the binding pose accuracy decreased as the ligand size increased, our models performed better for larger ligands than the baseline methods (Figure 2E), indicating their ability to efficiently handle many degrees of freedom. Besides, our models were able to sample more accurate binding poses for complexes with more related training samples (Figure 2F). This indicates that the modeling accuracy is dependent on the training data, and suggests that it may be possible to improve performance by enriching the training data or addressing any biases.

**Figure 2:**
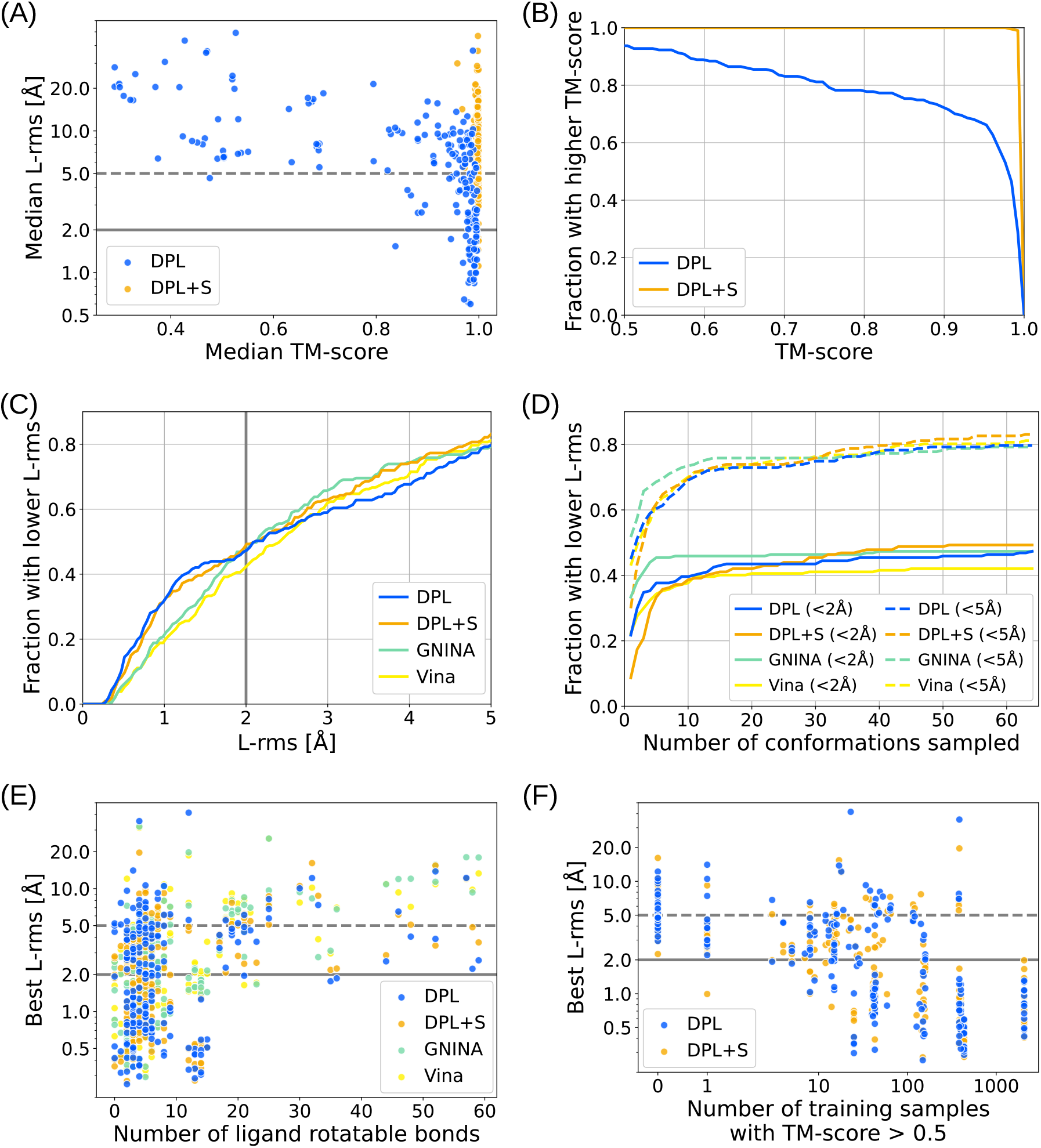
Benchmark results on the PDBbind test set. (A) Median modeling accuracies for each complex in the test set compared between our models DPL (blue) and DPL+S (orange). (B) Fraction of complexes where a structure with TM-score above the threshold, varying from 0.5 to 1.0, was sampled. (C) Fraction of complexes where a structure with L-rms below the threshold, varying from 0 Å to 5 Å, was sampled. The results obtained with the molecular docking methods (GNINA and AutoDock Vina) are also shown. (D) Performance as a function of the number of generative samples. Two thresholds 2 Å and 5 Å are employed for L-rms. (E) Relation between performance and the number of ligand rotatable bonds. (F) Relation between performance and the number of related training data.

### Effectiveness of protein structure-free modeling

In this experiment, we validated the effectiveness of protein structure-free modeling in a more challenging situation where ligand-bound protein structure was not available. We performed conformational sampling on 23 complexes from the PocketMiner dataset [43], a collection of apo-holo protein structure pairs with significant conformational changes upon ligand binding. These 23 complexes were selected by eliminating those with multiple annotated ligands annotated and those used for the training from the original dataset of 38 complexes. For methods that require protein structures as input, we examined both cases using the apo and holo structures.

Figure 3 shows the modeling accuracies on the PocketMiner dataset. As can be seen from Figures 3A and B, the performance of our models was not so good as for the PDBbind test set, and was worse than the docking with the holo structures. A possible reason for this might be that nearly half of the complexes (11 of the 23) in the PocketMiner dataset were out-of-distribution, i.e., no related samples (TM-score *>* 0.5) were observed during the training. Nevertheless, DPL outperformed both DPL+S and GNINA when the holo structures were unavailable (Figure 3B), as the performance of these structure-dependent methods was significantly degraded using apo structures (Figure 3C). In fact, DPL was able to sample structures with L-rms less than 5 Å on 10 of the 17 complexes where the docking with the apo structure failed. It is also worth noting that the superior performance of DPL was observed even on the out-of-distribution complexes (Figure 3C). This implies the generalization ability and applicability to complexes without related protein structures for training.

**Figure 3:**
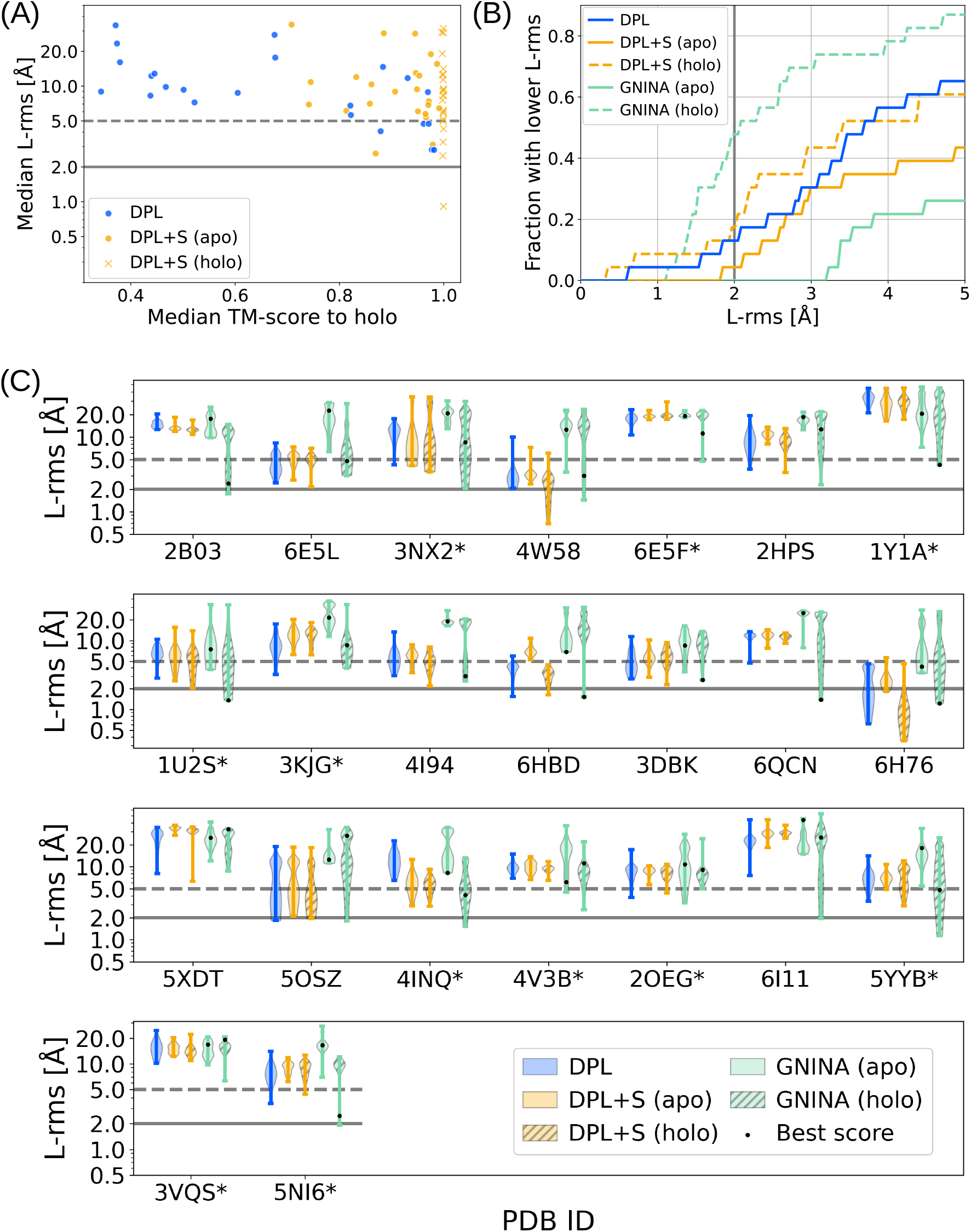
Modeling accuracies on the PocketMiner dataset. (A) Median modeling accuracies for each complex in the PocketMiner dataset, where the results by DPL, DPL+S (apo), and DPL+S (holo) are compared. (B) Fraction of complexes where a structure with L-rms below the threshold, varying from 0 Å to 5 Å, was sampled. The results obtained with GNINA are also shown for the apo and holo structures. (C) Distribution of L-rms for each complex in the PocketMiner dataset. The * on the PDB ID indicates that no related sample (TM-score *>* 0.5) was observed during the training.

An example of a Casein kinase II subunit alpha with an inhibitor bound to the substrate binding site (PDB ID 5OSZ) is presented in Figure 4. In this example, the correct binding pose could not be sampled by the docking with the apo structure because its binding site conformation was unsuitable for ligand binding (Figure 4C). In contrast, DPL successfully generated proper complex structures without using the holo structure (Figures 4A and D), demonstrating its effectiveness.

**Figure 4:**
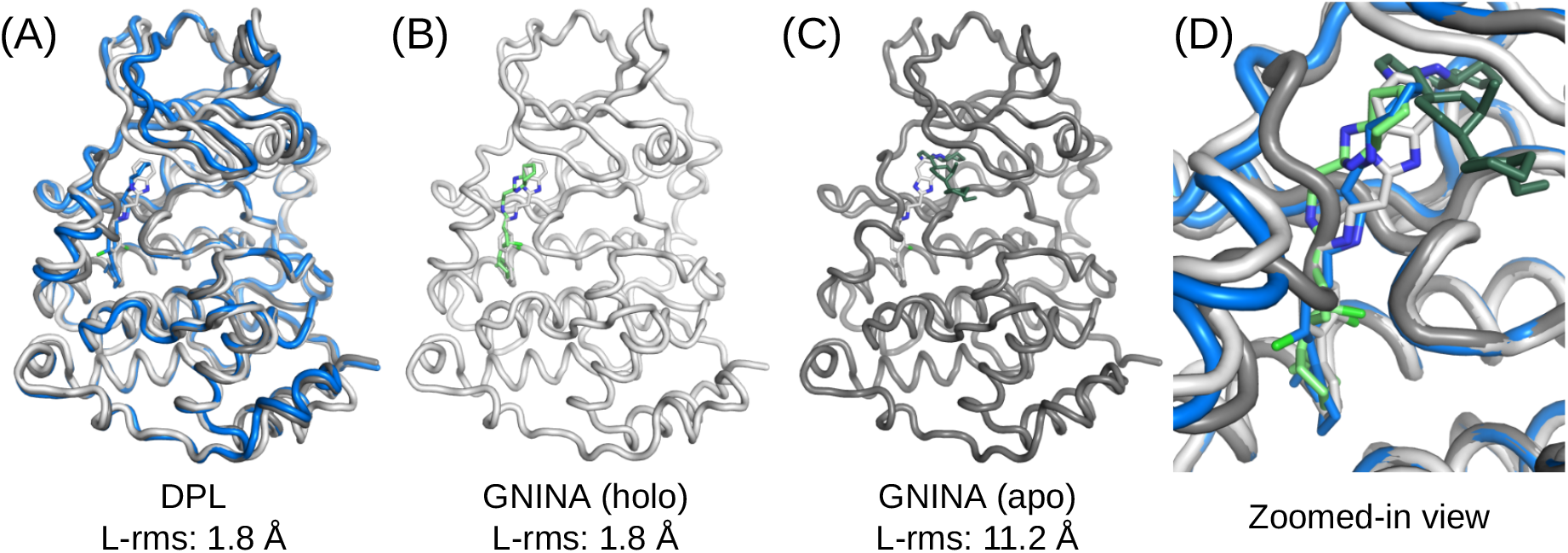
An example of a Casein kinase II subunit alpha with an inhibitor bound around the substrate binding site. The PDB-registered holo structure is shown in light gray (PDB ID: 5OSZ), and the apo structure is shown in dark gray (PDB ID: 6YPK). (A) Best L-rms complex structure generated by DPL (blue). (B) Best L-rms binding pose generated by GNINA using the holo structure (light green). (C) Best L-rms binding pose generated by GNINA using the apo structure (dark green). (D) Superimposition of all structures zoomed in on the ligand binding site.

Figure 5 illustrates the distribution of protein conformations generated by DPL for the complex of Sucrose-phosphatase and alpha-D-glucose (PDB ID: 1U2S). Even though no structure related to this complex was observed during the training, DPL was able to sample diverse protein conformations (Figure 5D), including both holo-like (Figure 5A) and apo-like (Figure 5C) ones. From the data in Figure 5D, we can also see that the input of protein structures biased the generated conformations so strongly that only small fluctuations were observed.

**Figure 5:**
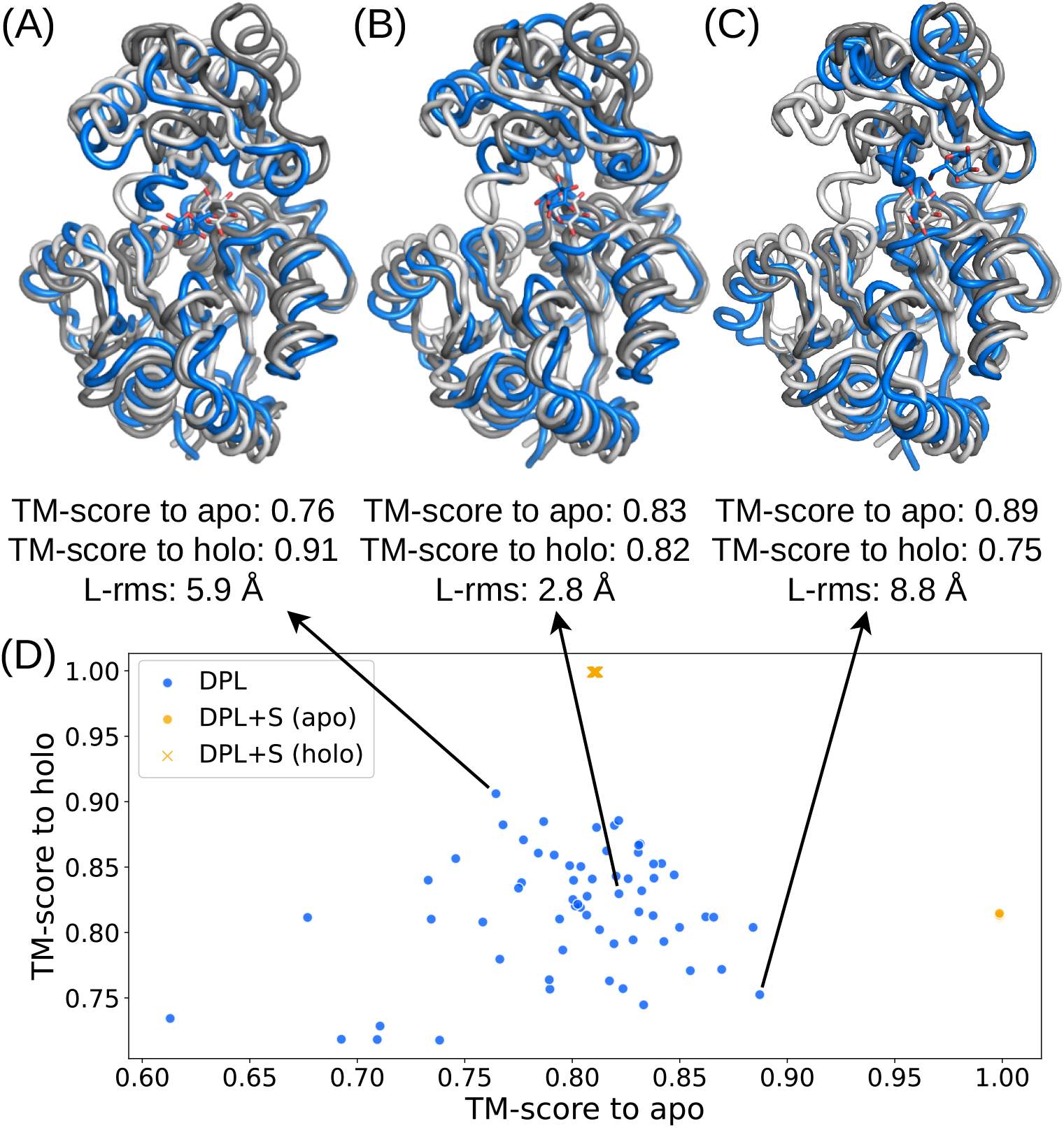
Distributions of protein conformations generated by DPL for the complex of Sucrose-phosphatase and alpha-D-glucose. The PDB-registered holo structure is shown in light gray (PDB ID: 1U2S), the apo structure is shown in dark gray (PDB ID: 1S2O), and the generated structure is shown in blue. (A) Generated structure closest to the holo structure. (B) Best L-rms generated structure. (C) Generated structure closest to the apo structure. (D) Distributions of the generated protein conformations projected onto the two-dimensional plane of TM-score to apo and TM-score to holo (blue symbols). The conformations obtained with DPL+S are also shown (orange symbols).

**Figure 6:**
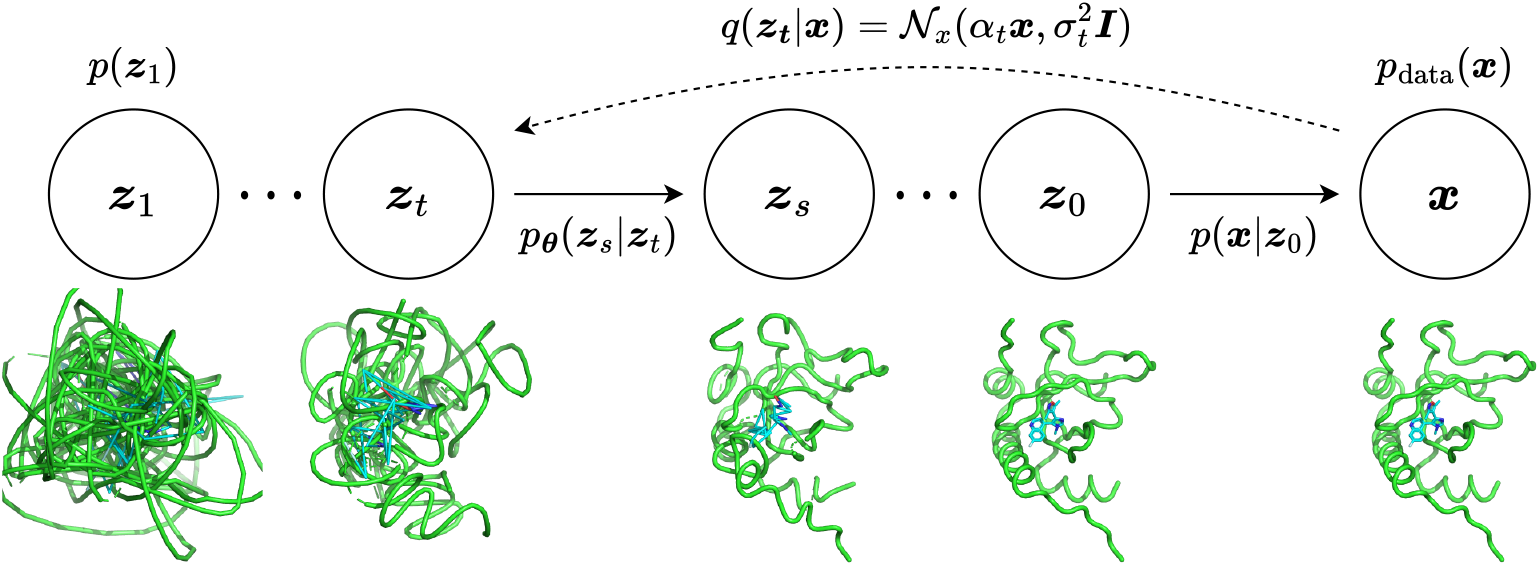
The diffusion and generative denoising process

## Discussion

The present results highlight the critical problem with previous protein structure-dependent approaches. Namely, preparing a protein structure suitable for ligand binding prior to determining the complex structure is an ill-posed problem, even though the sampling of binding poses is sensitive to inputs in these methods. Our approach solves this problem by generating the structure of protein-ligand complexes end-to-end without being biased by the input of protein structures.

Our protein structure-free model DPL demonstrated the ability to sample complex structures in many protein-ligand complexes with binding poses comparable to those obtained through molecular docking and showed generalization. However, its performance on complexes with little training data was still limited. This may be due, in part, to the bias in our training dataset, where some complexes had a thousand times more data than others, which could hinder generalization. In addition, the dataset size was small compared to the total number of entries in the PDB. Extensive use of PDB-registered structures, including apo protein structures, for training would improve the reproducibility of protein structures and help better capture the effects of ligand binding. Recent studies on protein structure prediction have reported improvements in prediction accuracy and generalization ability through the refinement of model architecture [17], training data [44], and scaling of pre-trained protein language models [18]. Based on these reports and our observations, we are optimistic that modeling accuracy and generalization ability will improve with better training data and the development of larger, more sophisticated models.

The protein conformational diversity observed in our experiments is remarkable, given that multistate sampling is a major challenge in current protein structure prediction methods. These methods are regression-based and rely on ad hoc techniques like MSA subsampling, changing the number of recycles, and enabling dropout during inference to sample multiple structures [44]. In contrast, our generative modeling approach was able to sample diverse protein structures in a principled way, without relying on these techniques. The effectiveness of generative models has also been discussed in the context of ligand docking [35].

Despite these promising results, our current method has some limitations. In particular, while we did not focus on scoring generated structures, this is crucial for practical applications like drug discovery. To address this issue, we could train a new scoring model as in DiffDock [35] or exploit likelihood estimation techniques using diffusion-based generative models [15, 29]. Besides, it is crucial to explicitly model all heavy atoms of the protein in order to interpret outcomes and combine them with other tools. Our method may be extended to the atomic level with little increase in computational cost by incorporating backbone orientations and side-chain dihedral angles in the diffusion model [32].

## Conclusion

We have introduced an equivariant diffusion-based generative model for end-to-end protein-ligand complex structure generation. The structures generated by our model were diverse and included those with proper protein conformations and ligand binding poses. When the ligand-bound protein structures were not available, our protein structure-free model showed better binding pose accuracy than the protein structure-dependent model and docking method, demonstrating the effectiveness of our end-to-end approach. The generalization observed in our experiments implies that our method can be applied to complexes with no related protein structures known, which has been challenging for earlier protein structure-dependent approaches. Based on these promising results, we conclude that the proposed framework will lead to better modeling of protein-ligand complexes, and we expect further improvements and wide applications.

## Methods

In this section, we describe the formulation of the equivariant diffusion-based model used in our framework, which is based on Variational Diffusion Models [26] and E(3) Equivariant Diffusion Models [29].

### The diffusion process

We first define an equivariant diffusion process for atom coordinates ***x*** and a sequence of increasingly noisy versions of ***x*** called latent variables ***z***_*t*_, where *t* ranges from *t* = 0 to *t* = 1. To ensure that the distributions are invariant to translation, we use distributions on the linear subspace [28] where the centroid of the molecular system is always at the origin and define 𝒩_*x*_ as a Gaussian distribution on this subspace, following [29]. The distribution of latent variable ***z***_*t*_ conditioned on ***x***, for any *t* ∈ [0, 1] is given by

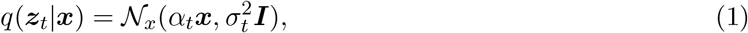

where *α*_*t*_ and 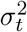 are strictly positive scalar-valued functions of *t* that control how much signal is retained and how much noise is added, respectively. We use the variance preserving process [23, 24] where 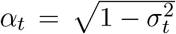, and assume that *α*_*t*_ is a smooth and monotonically decreasing function of *t* satisfying *α*_0_ ≈ 1 and *α*_1_ ≈ 0. Since this diffusion process is Markovian, it can also be written using transition distributions as

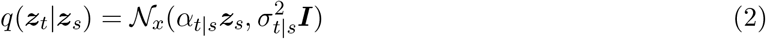

for any *t > s* with *α*_*t*|*s*_ = *α*_*t*_*/α*_*s*_ and 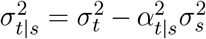. The posterior of the transitions given ***x*** is also Gaussian and can be obtained using Bayes’ rule:

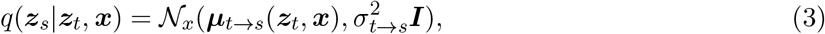

where

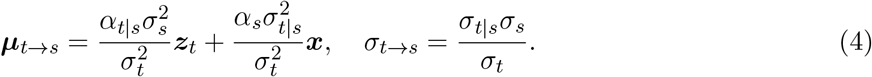

### The generative denoising process

The generative model is defined by the inverse of the diffusion process, where a sequence of latent variables ***z***_*t*_ is sampled backward in time from *t* = 1 to *t* = 0. Discretizing time uniformly into *T* timesteps, we can define the generative model as

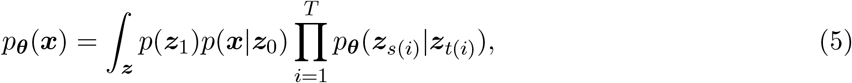

where *s*(*i*) = (*i*− 1)*/T* and *t*(*i*) = *i/T* . The variance preserving specification and the assumption that *α*_1_ ≈ 0 allow us to assume that *q*(***z***_1_) = 𝒩_*x*_(**0, *I***). We thus model the marginal distribution of ***z***_1_ as a standard Gaussian:

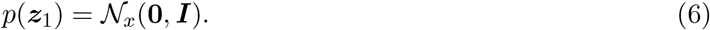

Similarly, with the variance preserving specification and the assumption that *α*_0_ ≈ 1, we can assume that *q*(***z***_0_|***x***) is a highly peaked distribution and *p*_data_(***x***) can be approximated as constant over this small peak. Therefore we have

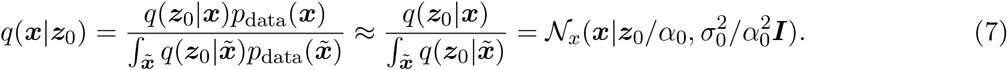

We then model *q*(***x***|***z***_0_) as

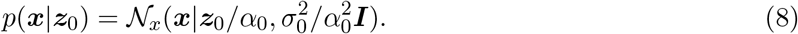

Finally, we define the conditional model distributions as

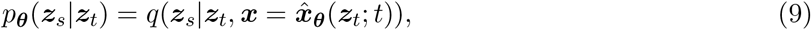

which is equivalent to *q*(***z***_*s*_|***z***_*t*_, ***x***), but with the original coordinates ***x*** being replaced by the output of a time-dependent denoising model 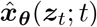 that predicts ***x*** from its noisy version ***z***_*t*_ using a neural network with parameter ***θ***. In practice, the denoising model is parametrized in terms of a noise prediction model 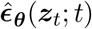:

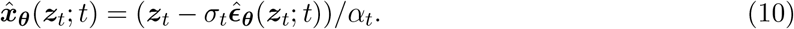

With this parameterization, 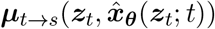 can be calculated as

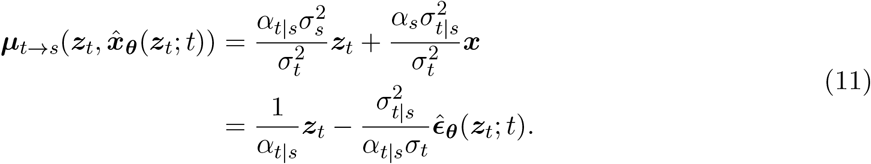

We can generate samples via ancestral sampling from this distribution (Algorithm 1).

#### Algorithm 1

Sample generation with noise prediction model 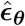.

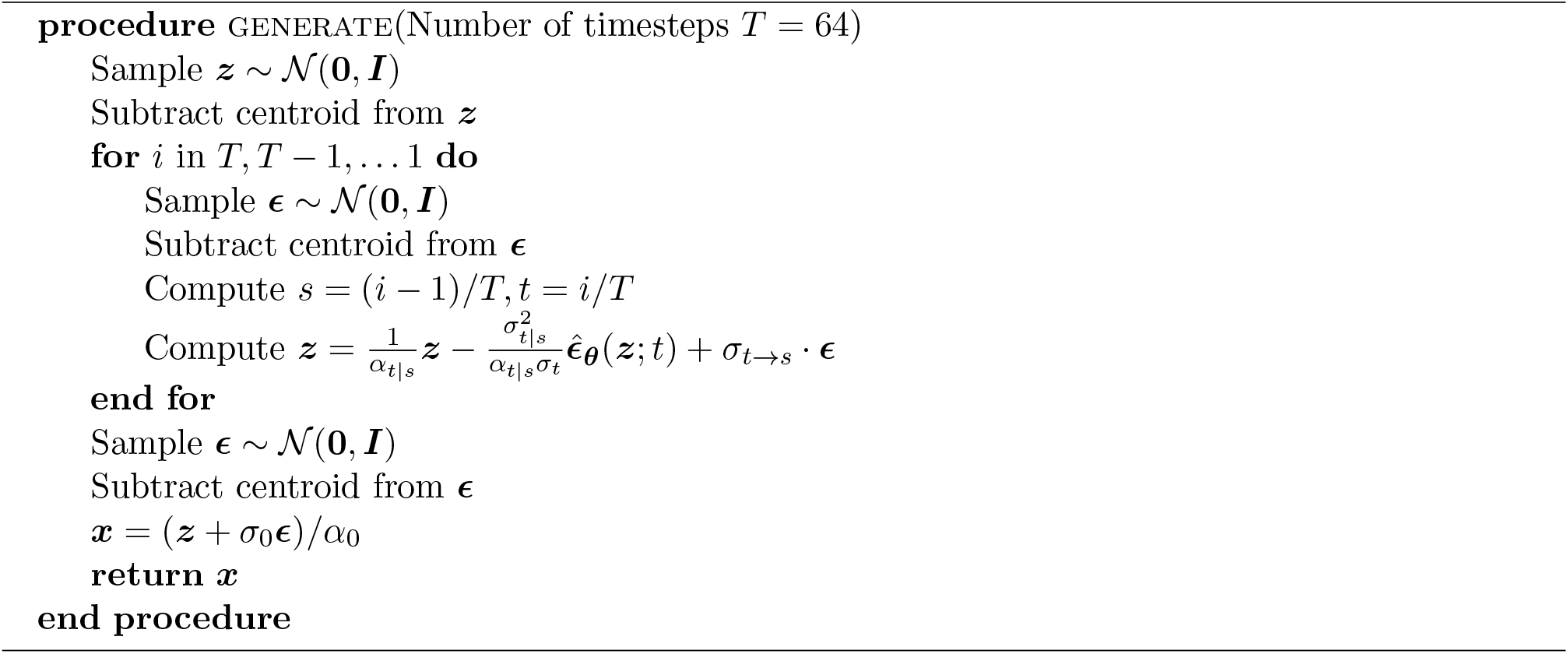

### Optimization objective

We optimize the parameters ***θ*** toward the variational lower bound (VLB) of the marginal likelihood, which is given by

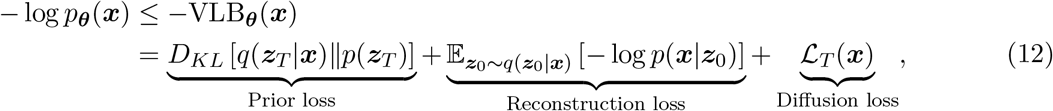

Where 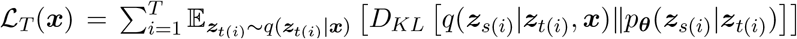. Since *p*(***z***_*T*_ ) and *p*(***z***_0_|***x***) contain no learnable parameter in our parameterization, the model is optimized by minimizing the third term, diffusion loss. As shown in [26], if we define the signal-to-noise ratio (SNR) at time *t* as 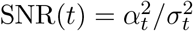 and *ϒ*(*t*) = − log SNR(*t*), then the diffusion loss can be simplified to

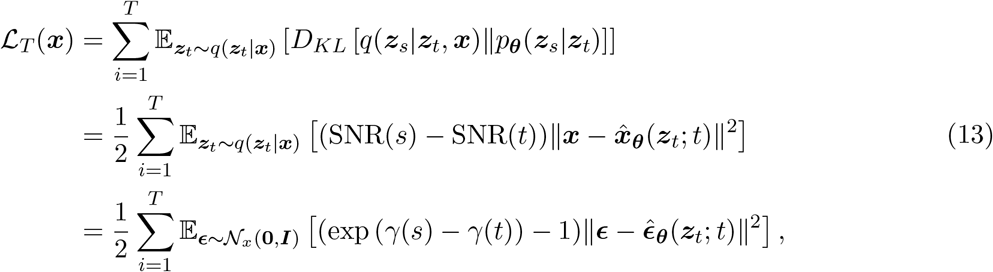

where *s* = (*i* − 1)*/T, t* = *i/T*, and ***z***_*t*_ = *α*_*t*_***x*** + *σ*_*t*_***ϵ***. Furthermore, we can consider a continuous time model corresponding to *T* → ∞. In the limit of *T* → ∞, the diffusion loss becomes

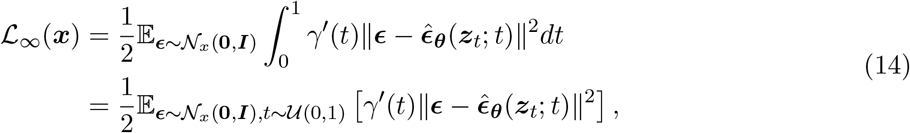

where *γ*^*′*^(*t*) = *dγ*(*t*)*/dt*. We use the Monte Carlo estimator of this continuous loss for parameter optimization (Algorithm 2).

### Model architecture

As illustrated in Figure 7, our noise prediction model consists of three procedures: (1) input featurization, (2) residual feature update, and (3) equivariant denoising. In this section, we outline each of these procedures, which are also described in Algorithm 3. For extensive details on the experimental setup, data, hyperparameters, and implementation, please refer to our code available at https://github.com/shuyana/DiffusionProteinLigand.

**Figure 7:**
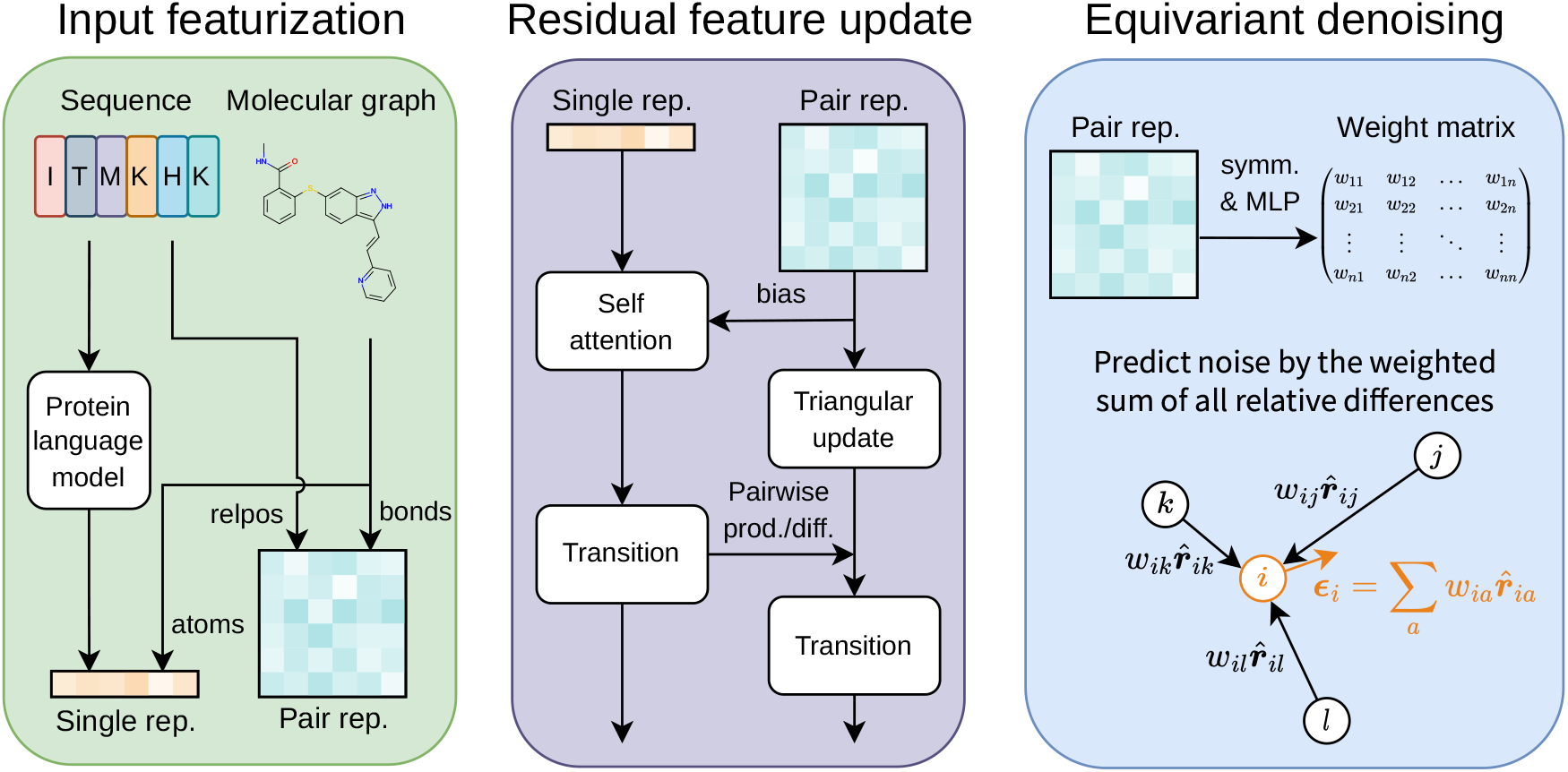
Overview of the model architecture. In the process of input featurization, single and pair representations are constructed. These features are then iteratively updated by the Folding blocks. The final pair representation is transformed by an MLP into a weight matrix to predict the denoising vector.

#### Algorithm 2

Parameter optimizaion of noise prediction model 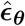.

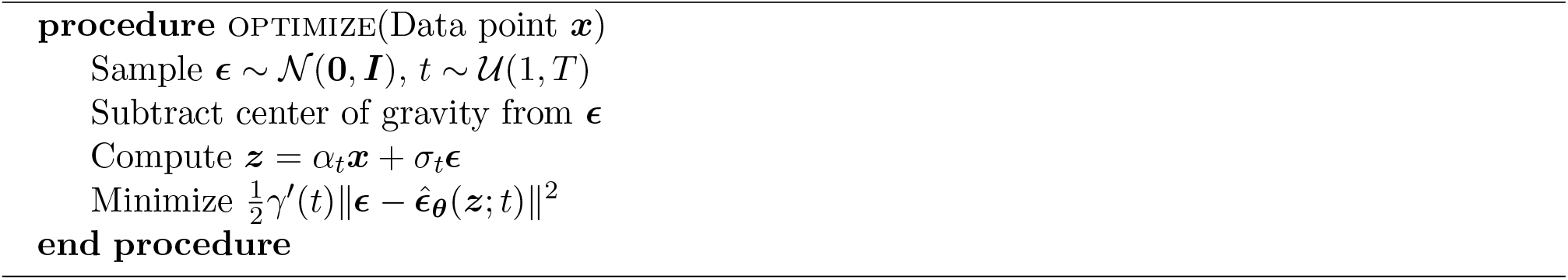

### Input featurization

We construct a single representation and a pair representation from a protein amino acid sequence and a ligand molecular graph. We utilize the 650M parameters ESM-2 [18] model, a large-scale protein language model pre-trained on ∼ 65 million unique protein sequences from the UniRef [45] database, to extract structural and phylogenetic information from amino acid sequences. To create the single representation of proteins, the final layer of the ESM-2 model is linearly mapped after normalization and then added to the amino acid embeddings. For the pair representation of proteins, we use the pairwise relative positional encoding described in the literature [17]. The representations of ligands are constructed through feature embedding of atoms and bonds. The features of the ligand atoms include: atomic number; chirality; degree; formal charge; the number of connected hydrogens; the number of radical electrons; hybridization type; whether or not it is aromatic; and whether or not it is in a ring. For ligand bonds, we use three features: bond type; stereo configuration; and whether or not it is considered to be conjugated. We concatenate the protein and ligand representations and then add them to the radial basis embeddings of atom distances and the sinusoidal embedding of diffusion time to obtain the initial representations of complexes.

### Residual feature update

We jointly update the single and pair representations with the 12 Folding blocks described in the ESMFold [18]. Although the Folding block was originally developed for proteins, it can also be used for protein-ligand systems without architectural modification.

### Equivariant denoising

In the process of equivariant denoising, the final pair representation is symmetrized and transformed by a multi-layer perceptron (MLP) into a weight matrix ***W*** . This matrix is used to compute the weighted sum of all relative differences in 3D space for each atom:

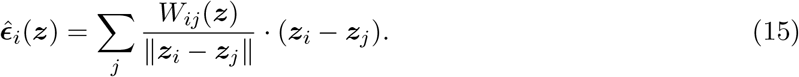

The centroid is then removed from this, resulting in the output of our noise prediction model 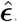.

Finally, we note that the model described above is SE(3)-equivariant, that is,

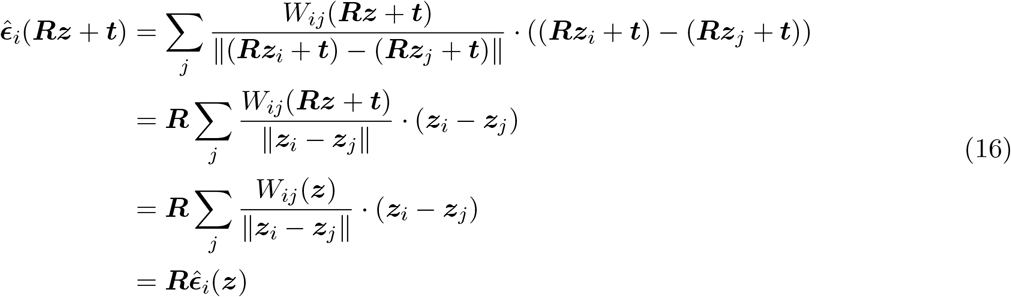

for any rotation ***R*** and translation ***t***. The second last equation holds because the final representation, and thus, the weight matrix ***W***, depend on atom coordinates only through atom distances that are invariant to rotation and translation.

## Acknowledgements

Not applicable.

## Abbreviations

MSA: multiple sequence alignment
PLM: protein language model
PDB: protein data bank
DPL: diffusion model for protein-ligand complexes
L-rms: ligand root mean square deviation
SNR: signal-to-noise ratio
MLP: multi-layer perceptron

## Algorithm 3

Noise prediction model. EmbedXXX embeds discrete features in representation space. ESM LastLayer returns the last hidden representations from an ESM language model.

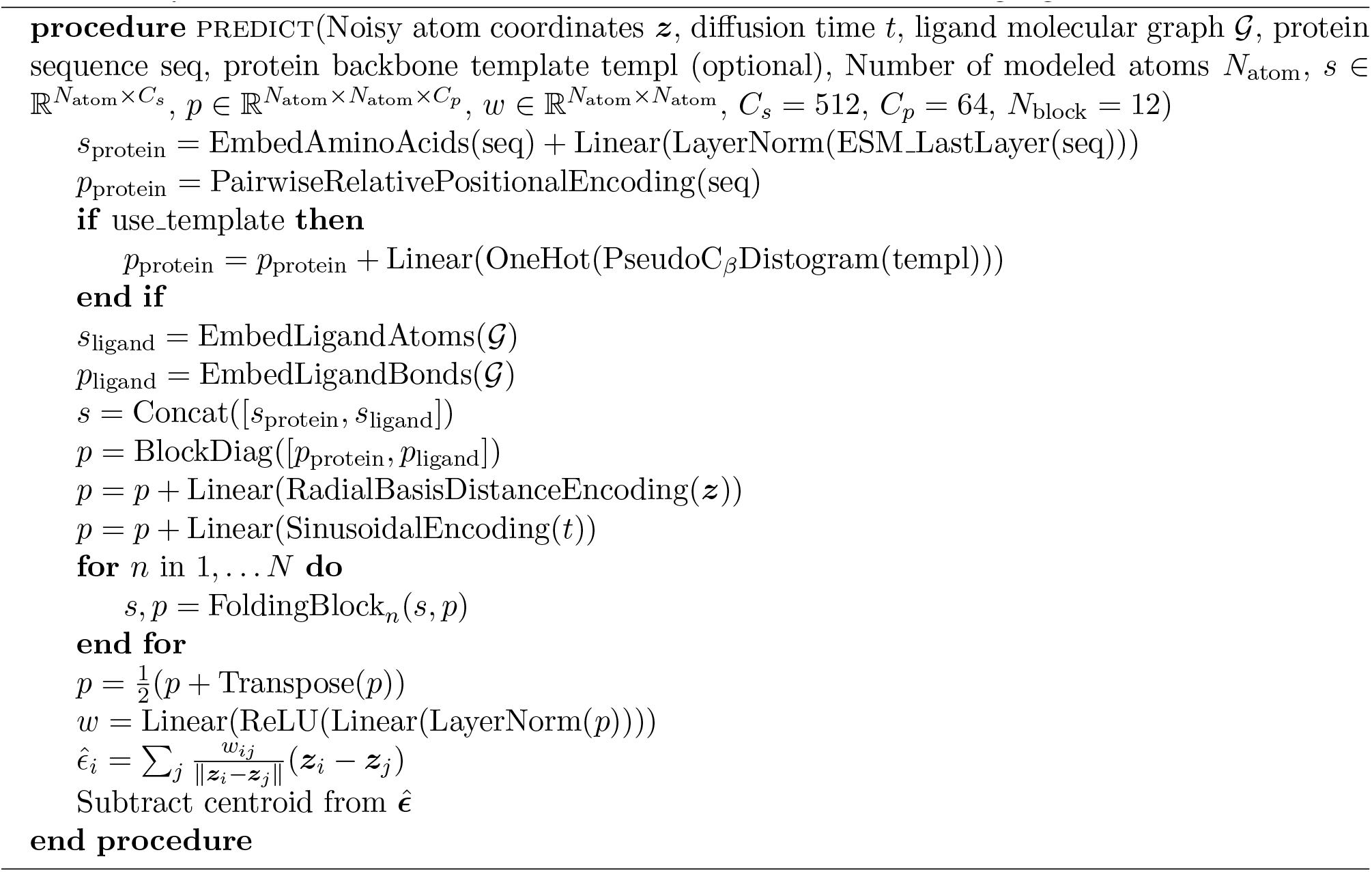

## Availability of data and materials

The PDBbind dataset is available at https://doi.org/10.5281/zenodo.6408497. The PocketMiner dataset is available at https://doi.org/10.1101/2022.06.28.497399. Our source code and model are available at https://github.com/shuyana/DiffusionProteinLigand.

## Ethics approval and consent to participate

Not applicable.

## Consent for publication

Not applicable.

## Authors’ contributions

SN analyzed and interpreted the numerical data in collaboration with YM and ST, and was a major contributor in writing the manuscript. All authors read and approved the final manuscript.

